# Extraction of near-complete genomes from metagenomic samples: a new service in PATRIC

**DOI:** 10.1101/2019.12.13.874651

**Authors:** Bruce Parrello, Rory Butler, Philippe Chlenski, Gordon D Pusch, Ross Overbeek

## Abstract

**Background:** Large volumes of metagenomic samples are being processed and submitted to PATRIC for analysis as reads or assembled contigs. Effective analysis of these samples requires solutions to a number of problems, including the binning of assembled, mixed, metagenomically-derived contigs into taxonomic units.

**Description:** The PATRIC metagenome binning service utilizes the PATRIC database to furnish a large, diverse set of reference genomes. Reference genomes are assigned based on the presence of single-copy universal marker proteins in the sample, and contigs are assigned to the bin corresponding to the most similar reference genome. Each set of binned contigs represents a draft genome that will be annotated by RASTtk in PATRIC. A structured-language binning report is provided containing quality measurements and taxonomic information about the contig bins.

**Conclusion:** We provide a new service for rapid and interpretable metagenomic contig binning and annotation in PATRIC.

## Background

### Context

Improvements in sequencing and assembly technology have created a wealth of metagenomic sample data. Because most organisms occurring in the world have not been successfully cultured in laboratory conditions, many species’ genomes are being reconstructed for the first time from metagenomic samples. This represents an important step towards a more complete characterization of true phylogenetic diversity and supports more accurate comparative analysis of microbial genomes.

In a recent paper, Pasolli *et al.* assembled over 153,000 high- and medium-quality draft genomes from metagenomic samples,[1] demonstrating the possibility of high-throughput metagenomic genome mining and indicating a future direction of data generation in genomics. It should be expected that, going forward, most new genomes being made available will be metagenomic in origin. The recently announced “Million Microbiome of Humans Project” suggests the scale of microbiome data that will become available in the near future.

In addition to genome discovery, rapid and accurate metagenomic sample characterization is in demand for a variety of industries including agriculture, ecology, and medicine. In particular, the ability to extract reasonably complete draft genomes from hospital samples without investing in specialized computing infrastructure is an important tool for diagnosis and research.

A key computational challenge in metagenomics is the binning of reads or assembled contigs into discrete taxonomic units. The high abundance of repeat regions, presence of DNA from low-abundance populations in the sample, inherent noisiness caused by the presence of multiple species in a single sample, and the lack of good reference genomes for unculturable populations make it challenging to produce full-length genomes from metagenomic reads.[2] Many of these problems affect metagenomic assembly, e.g. through assembly of chimeric contigs, and then propagate these problems into the subsequent binning step.[3] Tools such as METABAT[4] offer unsupervised binning of metagenomically-derived contigs and are considered state-of-the-art, but are computationally demanding and presently lack a stable online implementation as a service.

### Application in PATRIC

PATRIC, the Pathosystems Resource Integration Center, is a web-based service providing omics data and analysis tools that support the analysis of prokaryote genomes.[5] At the time of publication, it contains 324,422 annotated bacterial genomes and 4,807 annotated archaeal genomes. This population is expected to double in the next year. All of the genomes use RASTtk[6] to provide a single, controlled vocabulary for annotation, as well as a suite of quality metrics such as EvalG and EvalCon[7] for assessing draft genome quality. This controlled vocabulary enables us to make detailed comparative quality assessments among genomes. Such assessments are a crucial tool for the interpretation and evaluation of metagenome binning results.

Metagenomic analysis is in demand as a bioinformatics service, and most state-of-the-art services have high overheads in terms of configuration and investment in specialized computing resources. Few if any currently existing pipelines for metagenomic analysis are available in a convenient frontend for immediate use. We aim to provide PATRIC users with an easy-to-use, low-overhead, detailed, rapid, and structured metagenome binning service. Although it will likely be possible to achieve better performance on any given dataset using bespoke methods and specialized tools, our goal is to achieve near-state-of-the-art performance in as many cases as possible.

### Construction

### Overview

The PATRIC metagenome binning service takes metagenomic samples as assembled contigs or paired-end reads, extracts and annotates whole genomes, places them in the user’s workspace, and indexes them in the PATRIC database as private genomes. If reads are submitted, the metagenome binning service first assembles them into contigs using metaSPAdes.[8] Contigs are then binned using a novel supervised binning algorithm in which reference genomes are selected by searching for universal proteins in the sample. In this paper, a **reference genome** is a good-quality PATRIC genome that serves as a template for extracting draft genomes from a sample; a **universal protein** is a gene that is expected to occur exactly once in any prokaryote.

The PATRIC service uses the phenylalanine tRNA synthetase, alpha subunit (*pheS*) gene as its signature universal protein. This gene is long enough (209–405 amino acid residues in Bacteria and 293–652 in Archaea) to provide sufficient resolution for distinguishing between organisms at the species level but sufficiently conserved to provide reasonable closest-neighbor estimates for undersampled clades. Both of the above considerations are relevant to the selection of the reference genome, which is essential to the success of the supervised binning method. In PATRIC, *pheS* is the most abundant single-copy gene, so the set of potential reference genomes is slightly larger than for other potential universal proteins.

We use *pheS* rather than more conventional 16S rRNA-based methods because the faster molecular clock and single copy number make inference easier. Several studies have shown that universal genes achieve equal or better performance than 16S rRNA at distinguishing between populations.[9][10] In practice, *pheS* will likely underperform 16S rRNA for making inferences across large phylogenetic distances; since the binning service depends on identifying nearest neighbors only, this consideration should be largely immaterial.

It is unclear how much of an impact misidentifying the nearest reference genome should have on binning. It should be expected that, so long as the nearest reference for a given bin is sufficiently far away from the references for the other bins, even misidentification of the reference should minimally impact the quality of the binning. In any case, serious misidentifications of reference genomes should happen only in cases where closer references are unavailable in PATRIC. Since the clinical significance of undersampled clades is largely unknown, the selection of pheS for supervised binning is consistent with the priorities of the PATRIC project and user base. Moreover, the expansion of the PATRIC database to include more genomes from undersampled clades should substantially reduce the incidence of such cases over time.

This implementation leverages the PATRIC database structure for efficient and accurate draft genome reconstruction by accessing a large set of reference genomes and leveraging quality assessment tools like EvalG and EvalCon to verify draft genome quality. The binning service relies on a number of PATRIC-specific quality metrics (EvalG completeness, EvalG contamination, and EvalCon fine consistency) which we describe in a recent paper.[7]

### Dataset construction

We leverage the existing PATRIC dataset of roughly 330,000 public prokaryotic genomes to create a reference set of pheS sequences for the metagenomic binning service. We winnow the subset of 230,597 quality-controlled genomes to produce roughly 193,980 reference genomes meeting the following quality requirements:

1. Contamination score less than or equal to 10%
2. Fine consistency score greater than or equal to 87%
3. Completeness score greater than or equal to 80%
4. Exactly one *pheS* gene of appropriate length (209–405 amino acid residues for bacteria, 293–652 for archaea).

### Description of binning pipeline

#### Contig preprocessing

The contigs in the sample are first quality-controlled to eliminate long ambiguity runs. Any contig with more than 12 ‘N’ characters in a row, more than 50 ‘X’ characters in a row, or fewer than 400 total base pairs is discarded before the binning process begins.

#### Initialize bins

The binning service begins by identifying all occurrences of *pheS* in the sample by BLASTing the contigs in the sample against a small database of *pheS* sequences. Since this produces many low-quality hits by default, hits are discarded if they do not adequately cover the *pheS* sequence, or if the target contig has less than 4-fold average coverage, or if the target contig is less than 400 base pairs in length. Once the low-quality hits have been discarded, each remaining bin is identified as a provisional bin.

A bin is identified by its *pheS* sequence and contains a set of contigs. At the end of the initialization step, there are *N* bins containing exactly one contig, namely the contig containing the *pheS* role. Each of these bins is expected to correspond to a single species in the sample, although bins may be merged in subsequent steps if they turn out to be closely related.

#### Assign reference genomes

The metagenome binning service uses supervised binning methods to sort contigs into draft genomes. It compares contigs to the reference set of genomes in order to place them into bins. By contrast, unsupervised methods will use intrinsic characteristics such as GC content to group contigs together. (At this point, only the contigs containing *pheS* instances will have been binned.)

To assign reference genomes, the binning service compares the DNA sequences of the *pheS* instances in each bin to the reference set and chooses the closest match. Two bins are merged if they belong to the same species.

Next, sets of discriminating protein 12-mers are computed based on the reference genomes. A sequence of 12 amino acids in a given genome is said to be a discriminating protein 12-mer of that bin if it occurs in that bin’s reference genome(s) and does not occur in any of the other reference bins. Discriminating protein 12-mers are specific to the given sample.

#### Bin contigs

A contig *C* is placed into the bin belonging to reference genome set *G* if *C* has at least 10 discriminating protein 12-mers in common with *G*, and no other reference genome has more discriminating protein 12-mers in common with *C*. Contigs failing to meet the minimum similarity of 10 discriminating protein 12-mers with any of the reference genomes do not get binned. This is done for each contig until all contigs are either binned or discarded.

Finally, the discarded contigs are checked for long DNA sequences (50 base pairs or more) exactly matching a successfully binned contig from the previous step. If there is such a match between a discarded contig and a binned genome, the discarded contig is also placed into the corresponding bin. This will bin an additional 10–20% of the contigs. Discarded contigs without such a match are never binned.

#### Evaluate bin quality

Each bin created in the previous step represents a draft genome and can be annotated using the RASTtk pipeline. These annotations are used to compute three metrics of quality: completeness, contamination, and fine consistency. In addition to the quality metrics, each genome produced by the binning service includes a structured-language report on the absences or occurrences of problematic roles, i.e. roles which do not occur the expected number of times in a given genome. The quality metrics and problematic role report provide an independent check on the quality of the metagenome binning and allows users to review potential errors.

A good bin is defined in the same way that reference genomes are selected for the initial search, that is: contamination *<* 10%, fine consistency *≥* 87%, completeness *≥* 80%, and a single copy of the *pheS* gene of appropriate length.

#### Postprocessing

For low-quality genomes, a final postprocessing step is applied wherein contigs that do not get annotated with any good roles during the annotation step are also discarded from the draft genome. In this case, a contig having no good roles means that, of all of the annotated roles represented on that contig, the multiplicity predicted by EvalCon does not match the observed multiplicity of roles across the entire genome bin. When more occurrences of a role are observed in a given genome bin than predicted by EvalCon, the occurrences with the highest protein similarity to each of occurence in the reference genomes are considered good. The remaining occurrences are considered contamination. Contigs with no good roles tend to be short or misassembled, and discarding these contigs helps ensure that draft genomes correspond to general expectations about gene content in a given organism.

## Utility

### Integration into PATRIC

The metagenome binning pipeline is integrated into the PATRIC website as a service. Users can upload assembled contigs or paired reads (in which case the reads are first assembled using metaSPAdes). The user must also specify an output folder and a name for this PATRIC job. No additional input is required. The binning service is designed to run with minimal user input and still provide state-of-the-art metagenome binning.

When completed, the metagenome binning job produces a job directory containing BinningReport.html, a structured-language HTML report on binning effectiveness, as well as FASTA files and annotation reports for each of the bins. Each bin spawns a separate annotation job, which can be viewed in the same way as a regular annotation job submitted through the PATRIC annotation service.

The binning report contains information about the input file and a table of quality metrics (reference genome, coarse consistency, fine consistency, contamination, completeness, contig count, DNA size, contigs N50, mean coverage, good *pheS*) and a link to the problematic role report. Any quality metrics falling below the good genome cutoff are highlighted in yellow. Additional information on the use of this tool through the PATRIC website can be found in the metagenome binning service user guide and tutorial, both of which are linked on the upload page and accessible via the help page.

## Discussion

### Effect of postprocessing

When run on a set of 639 metagenomic samples from SRA, the pipeline without the postprocessing step produces an average of 1.7 good genome bins per metagenomic sample. After postprocessing, good genome yield increased to 8.17 good genome bins per sample.

### Effect of universal role selection

For the same set of 639 samples, we compared bins computed using Phenylalanyl-tRNA synthetase alpha chain (PheS) and N(6)-L-threonylcarbamoyladenine synthase (NLThreSynt). PheS failed to find any bins in 11 samples. NLThreSynt failed to find any bins in 14 samples. 8 of these failed instances were in common, so there are 3 samples for which PheS failed but NLThreSynt worked, and 6 samples in which NLThreSynt failed but PheS worked.

In 122 samples, NLThreSynt found more good bins than PheS. In 267 samples, PheS found more good bins. In 305 samples, NLThreSynt found more total bins, while PheS only found more total bins in 141 samples. NLThreSynt found 11,778 total bins and 4,715 good bins. PheS found only 11,180 total bins, but 5,163 good bins. Thus NLThreSynt’s production per sample is 7.37 good bins and 18.43 total bins; PheS’s production per sample is 8.08 good bins and 17.50 total bins.

### Comparison to other binning methods

In comparison with a recent study by Pasolli *et al.* featuring a custom unsupervised binning pipeline, the PATRIC metagenome binning pipeline achieved comparable results when run on a random set of 23 samples used in their study. In total, Pasolli *et al.* found 387 genome bins, of which 210 are good according to our definition. We found 370 bins, of which 206 were good. Since the Pasolli *et al.* study discovered a wide range of previously undiscovered and unculturable species, their use of unsupervised binning gives them a major advantage in this comparison. We hope that integrating their data into the PATRIC database will result in better performance in undersampled clades.

### Limitations and impact of new data

Because this service uses a supervised method, its effectiveness is highly dependent on the availability of high-quality reference genomes. Because the effectiveness of the binning service hinges on a binned genome being closer to its reference than to the references of all the other binned genomes, rather than simply on raw similarity, it is hard to characterize effect *a priori*. Nevertheless, the method is expected to underperform in undersampled clades.

Many metagenomically derived undersampled clades are undersampled because they resist cultivation *in vitro*. Thus, reference genomes must be produced on the basis of metagenomic samples.

On the one hand, increased attention to assembling and binning genomes from metagenomic samples, and in particular unsupervised binning, helps to characterize the diversity of metagenomic populations and fill in gaps in phylogenetic space. We expect the effectiveness of our supervised binning service to increase with the introduction of new references derived from unsupervised binning studies like Pasolli *et al*.

On the other hand, the lack of ground truth for metagenomically derived genomes is concerning. In particular, including metagenomically-derived genomes as references can proliferate errors in metagenomic assembly, binning, and annotation arbitrarily deep into undersampled clades. Chimeric assemblies can become established in the database as legitimate organisms, which would seriously affect the contig binning process. The metagenome binning service does not make users’ genomes public, and therefore they are never used as reference genomes, avoiding a feedback loop. Currently, PATRIC’s quality metrics control the quality of reference genomes derived from other sources in order to avoid propagating assembly and binning errors into the binning pipeline, but further checks on metagenomic draft genomes may be needed.

### Future improvements

In addition to the failsafes described above, PATRIC currently lacks a framework for dealing with metagenomically-derived bins or contigs as an aggregate, e.g. for the purpose of metatranscriptomics. The metagenome binning service currently creates a series of single-bin annotations and a genome group which supports some rudimentary multi-genome functionality, e.g. protein family heatmaps. Additionally, the current PATRIC metagenome binning service does not have tools for handling the presence of plasmids, phage and other viruses, or single-celled eukaryotes. These all represent potential areas for improvement of PATRIC’s support for metagenomics research.

## Conclusions

We have presented a service that takes paired-end reads or assembled contigs and produces a set of high-quality RASTtk-annotated contig bins. It additionally produces a report on the quality of all genome bins produced. This tool can be accessed via the PATRIC website through the Metagenomic Binning Service.

**Table 1.**
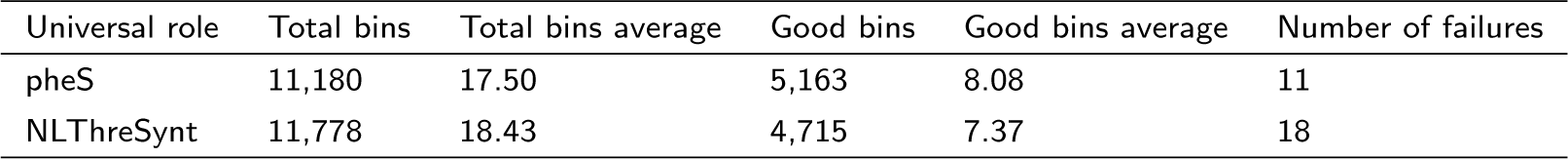
Comparison of universal role performance. Comparison of universal role performance on a random subset of 639 samples from the Pasolli et al study. In theory, this binning algorithm can be used for any single-copy universal marker gene. However, because pheS is the most abundant universal marker gene in PATRIC in practice (i.e. is missing from the fewest samples), it attains marginally better performance than other marker genes.

| Universal role | Total bins | Total bins average | Good bins | Good bins average | Number of failures |
| --- | --- | --- | --- | --- | --- |
| pheS | 11,180 | 17.50 | 5,163 | 8.08 | 11 |
| NLThreSynt | 11,778 | 18.43 | 4,715 | 7.37 | 18 |

**Figure 1.**
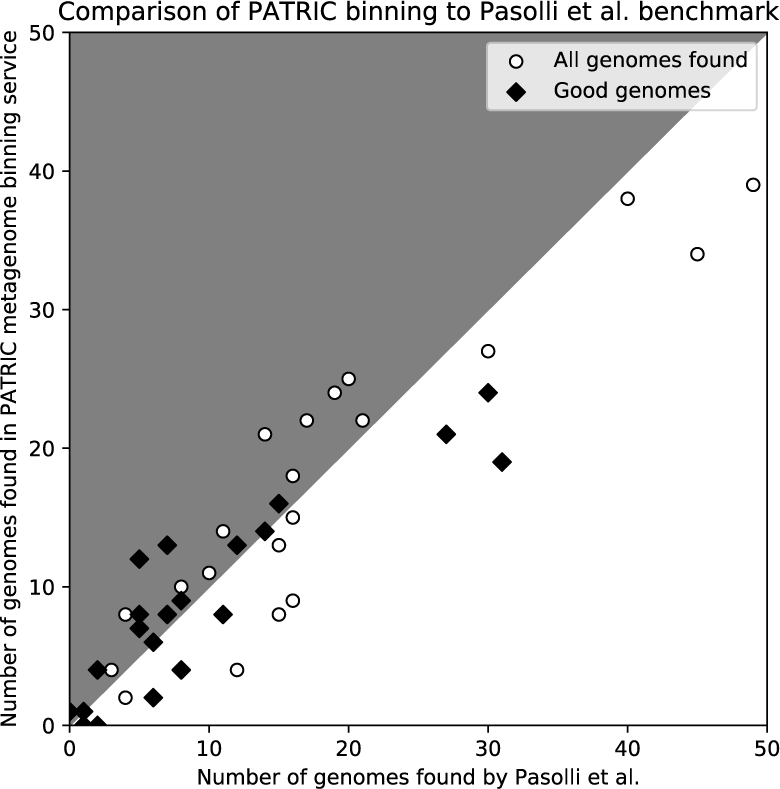
Performance on samples from Pasolli et al. study. This plot indicates the number of total bins and “good” bins (according to our criteria) generated by Pasolli *et al.* in their study and by the PATRIC metagenome binning method. Each point represents a sample, and each sample appears twice in the plot: once as the total number of bins found, once as the number of good bins only. Since Pasolli *et al.* is a good approximation of the state of the art in metagenome binning, this test suggests that the PATRIC binning method achieves performance close to the state of the art.

**Figure 2.**
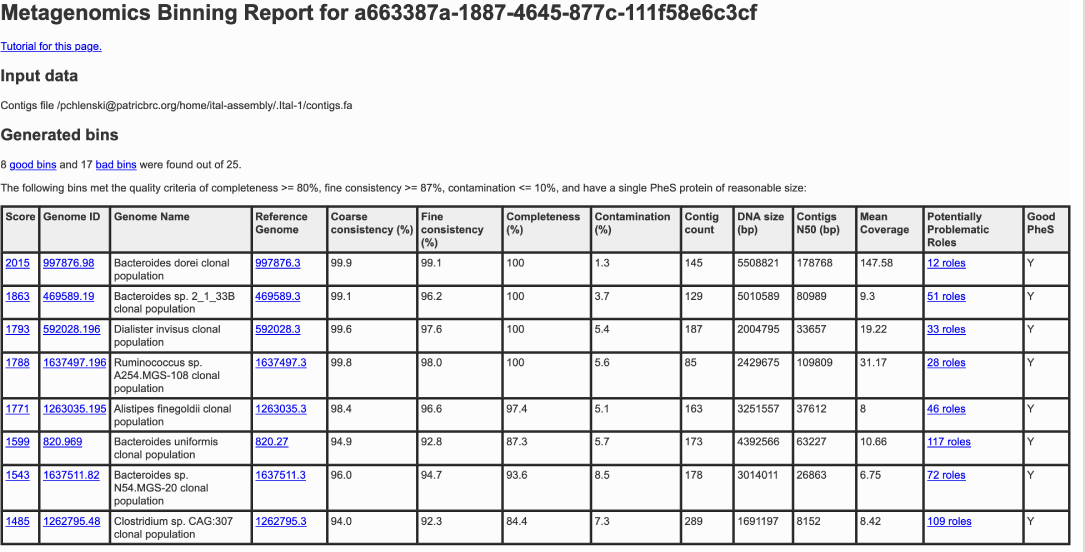
Binning report for good bins. The first part of the generated binning report summarizes the inputs, total number of bins, and quality breakdown of the bins. For good bins, it reports the overall score, genome ID (with link to PATRIC genome page), organism name (ending in “clonal population”), a link to the reference genome used in the binning process, and a number of quality metrics including a link to the problematic roles report.

**Figure 3.**
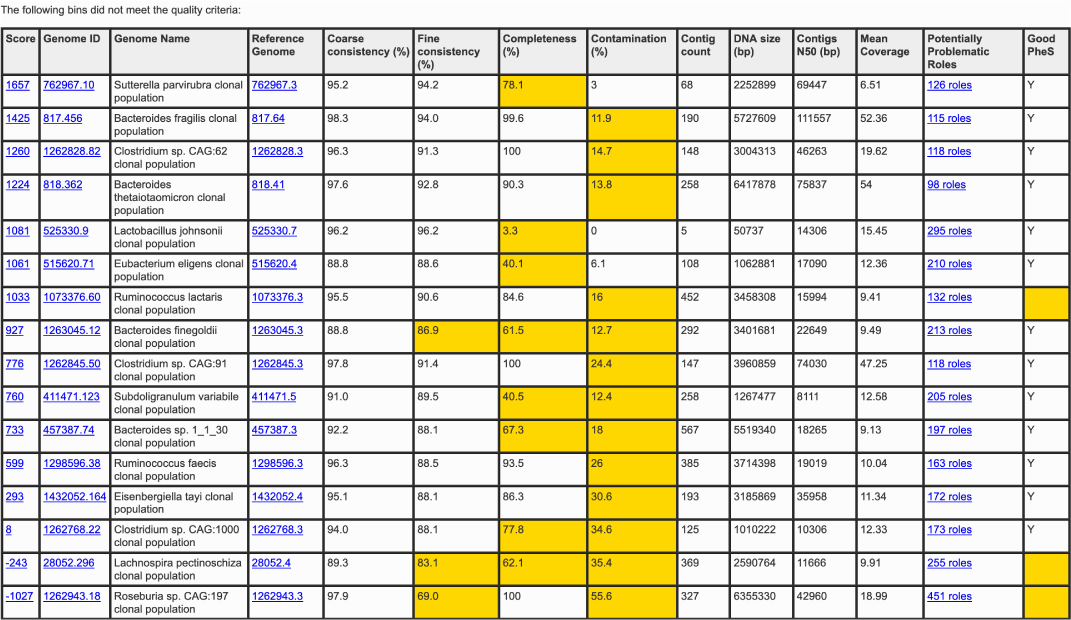
Binning report for other bins. The second part of the generated binning report lists the low-quality bins found. The columns are the same as for the good bins, but any metrics that fail to meet the criteria for a good bin are highlighted in yellow. This gives researchers a starting point into identifying what may have gone wrong in the assembly, binning, and/or annotation process.

## List of abbreviations

PATRIC: Pathosystems Resource Integration Center
RAST: Rapid Annotation Using Subsystems Technology
pheS: Phenylalanine tRNA synthetase, alpha subunit
SRA: Sequence Read Archive
NLThreSynt: N(6)-L-threonylcarbamoyladenine synthase

## Ethics approval and consent to participate

Not applicable.

## Consent for publication

Not applicable.

## Availability of data and materials

EvalCon and EvalG are available as part of the PATRIC annotation service, which can be accessed at https://patricbrc.org/app/MetagenomicBinning

## 1 Authors’ contributions

RO and BP developed the binning algorithm. BP developed the implementation of the algorithm. GDP, PC, and RB tested, benchmarked, and documented the binning pipeline performance, contributing to development of postprocessing methods. This manuscript was prepared by PC in consultation with the rest of the authors. All authors read and approved the final manuscript.

## 2 Acknowledgments

This material was based upon work supported by the U.S. Department of Energy, Office of Science, under contract DE-AC02-06CH11357. We thank Ralph Butler for his contributions to this paper.

